# Design of a holder for the improved maneuvering of the concurrent TMS-MRI setup

**DOI:** 10.1101/2023.03.12.530596

**Authors:** Hsin-Ju Lee, KJ Woudsma, Moh Ishraq, Fa-Hsuan Lin

## Abstract

**Background:** Concurrent transcranial magnetic stimulation (TMS) and magnetic resonance imaging (MRI) is time-consuming because of the limited space in the MRI bore and the sophisticated placement and orientation of the TMS coil to elicit the desired brain activities and behaviors.

**Objective:** We developed a TMS coil holder capable of quick adjustment of the TMS coil position and orientation. The holder can also hold an MRI receiver coil array.

**Methods:** A holder with one controlling knob, two omni-direction rotation joints, and two in-plane rotation joints was developed.

**Results:** Different TMS coil positions and orientations can be arranged and fixed in seconds. The holder can also accommodate two TMS coils to allow for multi-coil TMS-MRI.

**Conclusion:** Our development significantly improves the workflow of the concurrent TMS-MRI in new neuroscience studies and clinical applications.

## Introduction

The combination of TMS and MRI allows for 1) imaging-guided neuro-intervention [1] and 2) immediate assessment of the stimulation effect [2,3]. For example, TMS at the bilateral horizontal segment of the intraparietal sulcus (hIPS) guided by a group-level task-fMRI revealed that the hIPS is required in arithmetic processing [4]. The frontal eye fields (FEF) revealed by an individual’s fMRI guided the rTMS and caused reduced alpha oscillations at the contralateral (to the stimulation site) occipitoparietal lobe, suggesting that FEF are causally involved in the topdown attentional control of anticipatory alpha oscillation of the visual system [5]. Clinically, personalized fMRI-guided TMS using resting-state fMRI for MDD treatment showed improved responses [6].

Practically, TMS inside the MRI bore is difficult because of the limited bore size. The TMS coil position and orientation may be first adjusted when a participant is outside the MRI or in a sit-up position to allow for more room to maneuver. As the subject goes inside the MRI bore, the found TMS coil location and orientation may be lost. Re-adjustment can be tedious because of separate buttons and handles controlling independent TMS coil rotation and translation in independent directions [7–9].

Multi-site TMS offers the possibility of modulating the brain function in a network manner: magnetic pulses can be delivered with controlled latencies at different target loci [10,11]. However, realizing multi-site TMS inside the MRI bore is more challenging in space management.

Here we present a novel design of the TMS coil holder, which can be easily controlled for high degrees of freedom with a single controlling knob. The coil holder also allows for positioning and orienting with in-plane rotation of the TMS coil. A setup is shown to allow for current TMS-MRI at a 3T MRI scanner and an 8-channel receiver coil array. The TMS coil holder is compact to allow for more than one TMS coil inside the bore. This design facilitates the development of TMS-MRI discovery and promotes the reproducibility of TMS-MRI findings.

## Material and methods

The design of the TMS coil holder is illustrated in **Figure 1A**. The holder consisted of three in-plane rotation (IPR) joints, two arms between two IPR joints, and two omnidirectional rotation (ODR) ball-head joints at both ends. Two ODR joints enabled 360^O^ rotation between the arm and end plate and the TMS coil. The controlling knob had a mechanism allowing for the simultaneous fixing and releasing of two ODR joints and one IPR joint at the center. The IPR joint at the TMS coil end allowed for in-plane rotation of the TMS coil. The base had multiple fixing points to accommodate more than one set of the TMS coil holder. Thus, a multi-coil TMS-MRI setup is possible. The rails for the base were designed to fix the holder on the MRI bed (Prisma, Siemens, Germany).

**Figure 1.**
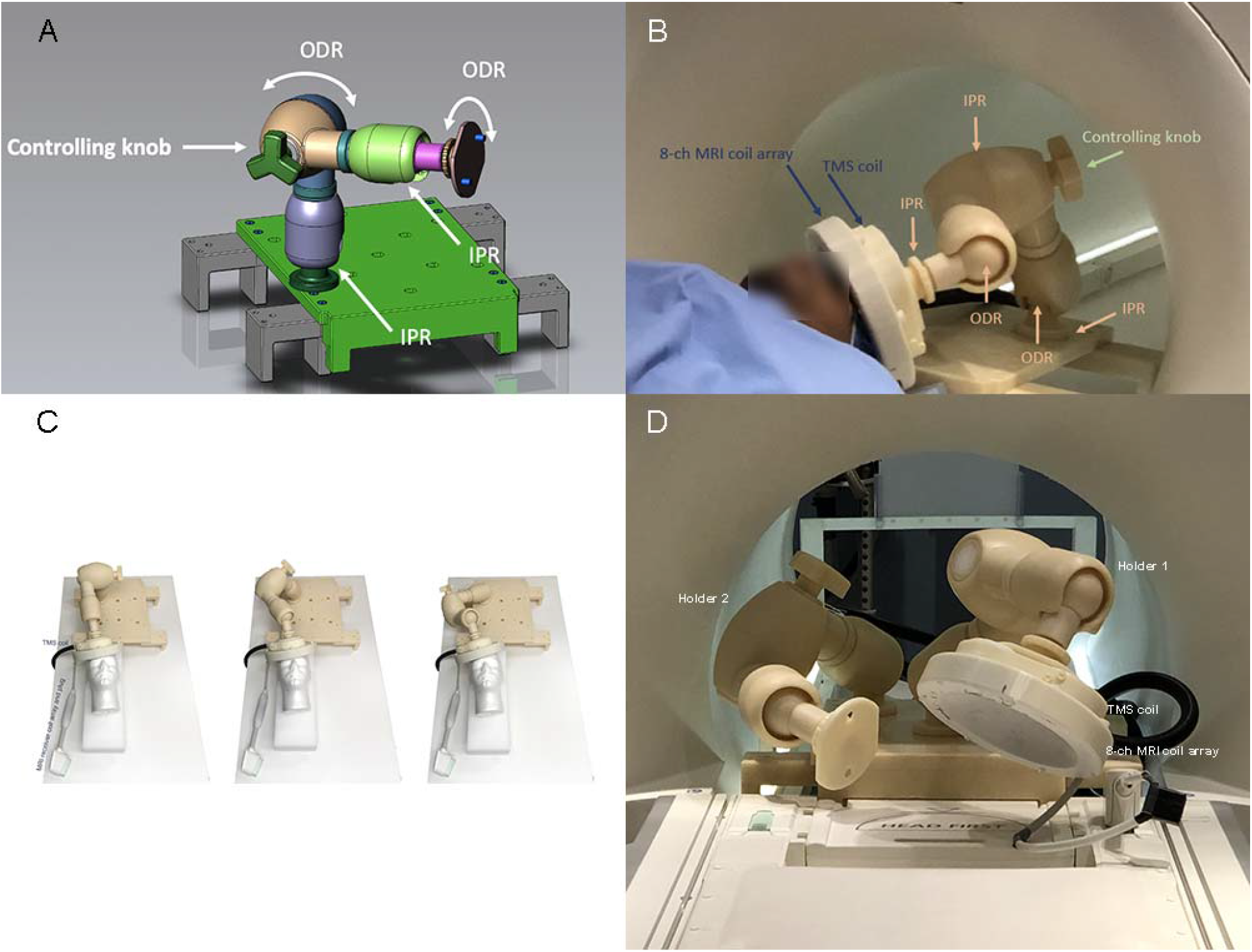
A. The design of the TMS coil holder for TMS-MRI experiments. B. The experimental setup of TMS-MRI with an MRI-compatible TMS coil, 8-channel MRI coil array, and coil holder. IPR: In-plane rotation joint. ODR: Omni-directional rotation joint. C. Three ways to place the TMS and MRI coils at the same locations and positions, illustrating the degree of freedom offered by the holder. D. An exemplary setup of two holders at the MRI. The empty Holder 2 can mount a TMS coil, MRI coil, supporting pad, or mirror.

The TMS coil was made of polycarbonate (Samferson Ltd., Taipei, Taiwan) by 3D printing. All parts and fixtures in the holder were non-magnetic. The total weight of the TMS coil holder, including the base, was 22 Kg.

We tested the coil holder outside the MRI to demonstrate its flexibility in optimizing the TMS coil position and orientation. An MRI-compatible TMS coil (MRi-B91, MagVenture, Denmark) was attached to the coil holder for inside-MRI testing. A homemade 8-channel coil array was attached to the TMS coil (**Figure 1B**) to collect MRI at the vicinity of the TMS stimulation target. We acquired MRI with an echo-planar imaging (EPI) sequence (TR: 2000 ms; TE: 30 ms; flip angle: 70^o^; resolution: 3×3×3 mm^3^; image matrix: 64; 20 slices) with this setup.

## Results

**Figure 1C** shows the setup of the TMS coil with three different configurations of the coil holder. Note that these three configurations gave the same TMS coil position and orientation by releasing and fixing the only controlling knob. These configurations exemplified the flexibility to arrange TMS coil orientation and position with this holder. **Figure 1C** also shows the setup of the TMS coil holder with a TMS coil and an 8-channel MRI coil array inside the MRI bore. We also show that two TMS coil holders can be mounted on the same base, illustrating the possibility of multi-loci TMS using separate TMS coils. **Figure 1D** shows the setup at MRI with two holders. One holder had the TMS coil and an 8-channel MRI coil array. The other holder had an adaptor for a possible TMS coil, MRI coil, supporting pad, or mirror. **Figure 2** shows the MRI of a healthy subject. As the TMS and MRI coil can be moved rather freely inside the MRI by controlling one knob, we achieved both coil translation (about 25 mm) and rotation (about 15^o^) within 30 s.

**Figure 2.**
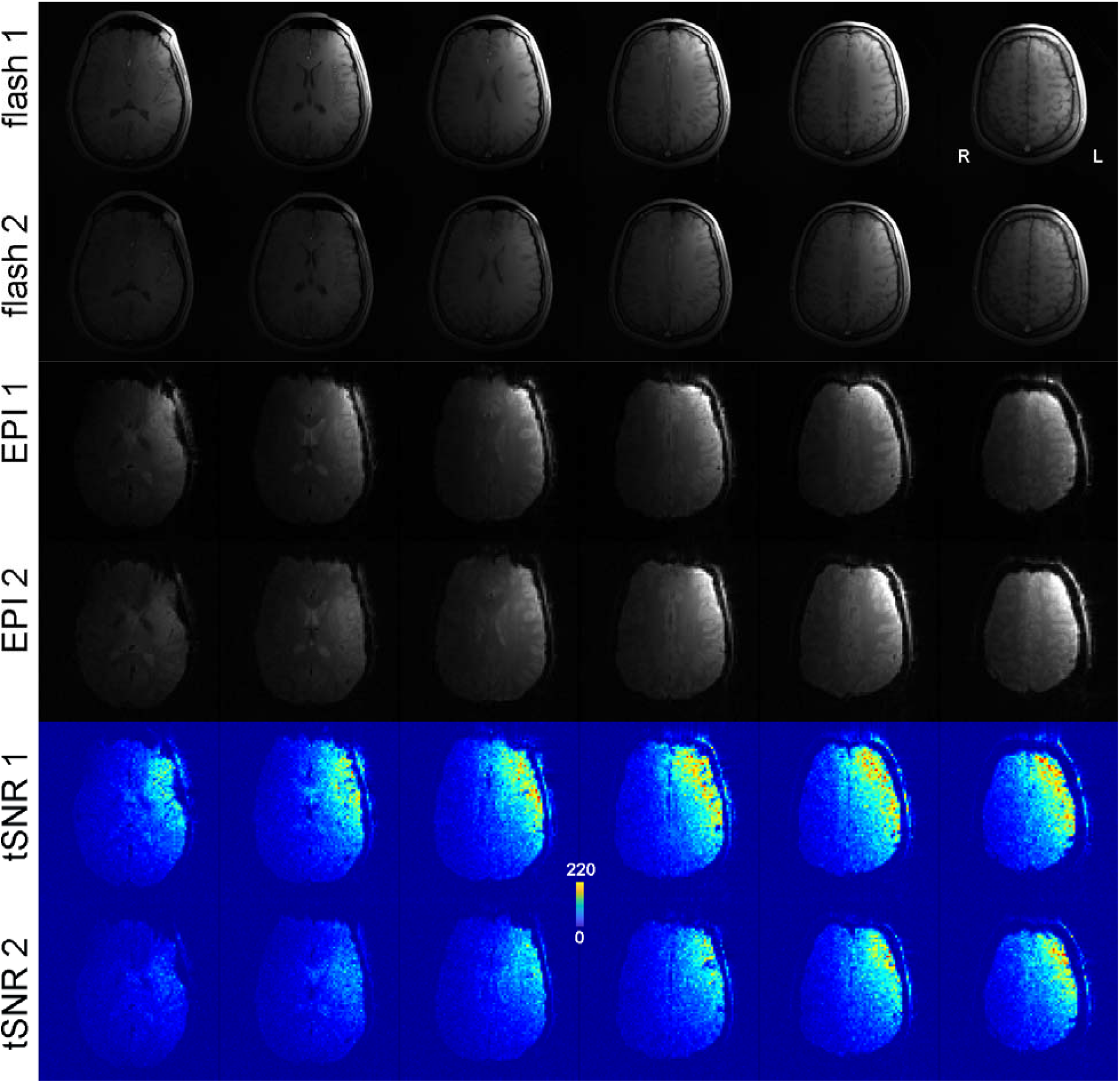
Top: Gradient-echo (FLASH sequence) and echo-planar imaging (EPI) images of the same participant with different MRI and TMS coil positions and orientations in two runs with the setup adjusted in 30 s. Bottom: time-domain signal-to-noise ratio (tSNR) maps of the EPI data in two runs. L: left. R: right.

## Discussion

A highly flexible TMS coil holder was developed for a concurrent TMS-MRI experiment. The mechanical design enabled fast (**Figures 1A** and **2**), a high degree of freedom (**Figure 1C**), and dedicated changes of the TMS coil within a limited space inside the MRI bore (**Figures 1B** and **2**). The current TMS coil holder needs to use separate knobs to control coil rotations and translations, which are in orthogonal directions. However, in practice, the research typically needs to move the TMS coil in non-orthogonal directions along a curved surface (i.e., scalp) in optimizing the TMS target. Our design greatly facilitated this procedure.

Note that our coil holders can hold more than one set of TMS and MRI coils (**Figure 1D**). Thus, multi-loci TMS can be more accessible. Importantly, our holder had different adaptors to attach to a TMS coil, an MRI coil array, and other mechanical supports. The application of this TMS coil holder was versatile. For example, an MRI coil array can be attached to a second TMS holder to enable imaging of brain activity outside the vicinity of the TMS target at the first TMS holder. Alternatively, the TMS holder can mount the mirror in functional MRI experiments, where visual stimuli can be shown to the participant to perceive the input from a projector at the mirror. A TMS coil holder can also serve as a mechanical support to reduce the participant’s motion when, for example, a TMS coil is supported by another TMS holder in the other hemisphere.

The TMS coil holder presented here needs to adjust its location and orientation manually. Constrained by the strong magnetic field inside the MRI system, conventional motors cannot be mounted on this holder for robotic control. However, it is likely to use MRI-compatible motors, such as hydraulic ones, to control ODR and IPR joints. This will open the possibility of robotic neurostimulation in the future.

## Acknowledgement

The authors thank the valuable discussion with Jason Shen from Samferson Ltd. Taipei, Taiwan, and Dr. Sean Nestor at Sunnybrook hospital.

## Disclosure of competing interests

All authors declare that there is no financial or personal relationship with other people or organizations that could inappropriately influence their work.

## Funding

This work was supported by the Natural Sciences and Engineering Research Council of Canada [grant number RGPIN-2020-05927]; Canada Foundation for Innovation [grant numbers 38913 and 41351]; Canadian Institutes of Health Research [grant number PJT 178345]; MITACS [grant number IT25405].

## Notes

### Competing Interest Statement

The authors have declared no competing interest.

## References

[1] Bergmann TO, Karabanov A, Hartwigsen G, Thielscher A, Siebner HR. Combining non-invasive transcranial brain stimulation with neuroimaging and electrophysiology: Current approaches and future perspectives. Neuroimage 2016;140:4–19.

[2] Peters JC, Reithler J, Graaf TA de, Schuhmann T, Goebel R, Sack AT. Concurrent human TMS-EEG-fMRI enables monitoring of oscillatory brain state-dependent gating of cortico-subcortical network activity. Commun Biol 2020;3:40.

[3] Bestmann S, Ruff CC, Blankenburg F, Weiskopf N, Driver J, Rothwell JC. Mapping causal interregional influences with concurrent TMS–fMRI. Exp Brain Res 2008;191:383–402.

[4] Andres M, Pelgrims B, Michaux N, Olivier E, Pesenti M. Role of distinct parietal areas in arithmetic: an fMRI-guided TMS study. Neuroimage 2011;54:3048–56.

[5] Marshall TR, O’Shea J, Jensen O, Bergmann TO. Frontal eye fields control attentional modulation of alpha and gamma oscillations in contralateral occipitoparietal cortex. J Neurosci 2015;35:1638–47.

[6] Cash RFH, Cocchi L, Lv J, Fitzgerald PB, Zalesky A. Functional Magnetic Resonance Imaging-Guided Personalization of Transcranial Magnetic Stimulation Treatment for Depression. JAMA Psychiatry 2021;78:337–9.

[7] Bohning DE, Denslow S, Bohning PA, Walker JA, George MS. A TMS coil positioning/holding system for MR image-guided TMS interleaved with fMRI. Clin Neurophysiol 2003;114:2210–9.

[8] Moisa M, Pohmann R, Ewald L, Thielscher A. New coil positioning method for interleaved transcranial magnetic stimulation (TMS)/functional MRI (fMRI) and its validation in a motor cortex study. J Magn Reson Imaging 2009;29:189–97.

[9] Goldstein S, Rafiei F, Rahnev D. 3D-printed stand, timing interface, and coil localization tools for concurrent TMS-fMRI experiments. Brain Stimul 2022;15:1290–1.

[10] Nieminen JO, Sinisalo H, Souza VH, Malmi M, Yuryev M, Tervo AE, et al. Multi-locus transcranial magnetic stimulation system for electronically targeted brain stimulation. Brain Stimul 2022;15:116–24.

[11] Ferreri F, Pasqualetti P, Määttä S, Ponzo D, Ferrarelli F, Tononi G, et al. Human brain connectivity during single and paired pulse transcranial magnetic stimulation. Neuroimage 2011;54:90–102.

